# Ermin deficiency as an inside-out model of inflammatory dysmyelination

**DOI:** 10.1101/2020.06.16.154781

**Authors:** Amin Ziaei, Marta Garcia-Miralles, Carola I. Radulescu, Harwin Sidik, Aymeric Silvin, Han-Gyu Bae, Carine Bonnard, Nur Amirah Binte Mohammad Yusof, Costanza Ferrari Bardile, Liang Juin Tan, Alvin Yu Jin Ng, Sumanty Tohari, Leila Dehghani, Byrappa Venkatesh, Sarah R. Langley, Vahid Shaygannejad, Bruno Reversade, Sangyong Jung, Florent Ginhoux, Mahmoud A. Pouladi

## Abstract

Ermin is an actin-binding protein found almost exclusively in the central nervous system (CNS) as a component of myelin sheaths. Although Ermin has been predicted to play a role in the formation and stability of myelin sheaths, this has not been directly examined *in vivo*. Here we show that Ermin is essential for myelin sheath integrity and normal saltatory conduction. Loss of Ermin in mice caused de-compacted and fragmented myelin sheaths and led to slower conduction along with progressive neurological deficits. RNA sequencing of the corpus callosum, the largest white matter structure in the CNS, pointed to inflammatory activation in aged Ermin-deficient mice, which was corroborated by increased levels of microgliosis and astrogliosis. The inflammatory milieu and myelin abnormalities were further associated with increased susceptibility to immune-mediated demyelination insult in Ermin knockout mice. Supporting a possible role of Ermin deficiency in inflammatory white matter disorders, a rare inactivating mutation in the *ERMN* gene was identified in multiple sclerosis patients. Our findings demonstrate a critical role for Ermin in maintaining myelin integrity. Given its near exclusive expression in myelinating oligodendrocytes, Ermin deficiency represents a compelling “inside-out” model of inflammatory dysmyelination and may offer a new paradigm for the development of myelin stability-targeted therapies.

## INTRODUCTION

Myelin is a multilayered, compact membrane structure that insulates and supports axons in the central (CNS) and peripheral (PNS) nervous systems^1^. The myelin sheath is exceptionally stable and is one of the most complex biological membranes, with unique composition and architecture. Despite its importance, much of the molecular and cellular choreography underlying the generation, assembly, maintenance, and remodeling of myelin sheaths and how it differs between the CNS and PNS remains poorly understood.

Although CNS and PNS myelin have similar functions, each is unique in several respects including their molecular composition^1^. For example, whereas proteolipid protein (PLP) is the most abundant protein constituent of myelin in the mammalian CNS, it represents less than 1% of PNS myelin. Conversely, the single membrane P0 glycoprotein, the most abundant protein in PNS myelin, is nearly absent from mammalian CNS myelin. The shift from P0 to PLP as a major constituent of CNS myelin is a key evolutionary event in the transition from fish to higher vertebrates2 and is paralleled by the emergence of a protein named Ermin expressed exclusively in the CNS and which localises to myelin sheaths of mature, myelinating oligodendrocytes^3–5^.

Unlike classical myelin proteins such as PLP, MBP, CNP, MOG and MAG, Ermin represents less than 1% of the myelin protein, making it a quantitatively minor myelin protein^6^. Ermin was first described as an oligodendrocyte-specific gene and a new member of the ERM (Ezrin, Radixin and Moesin) family of proteins^4, 5^. ERM proteins cross-link actin filaments with plasma membranes in diverse cell types, including Schwann cells^7^. Structurally, Ermin exhibits key similarities and differences with other ERM proteins: it shares a highly conserved C-terminus actin binding domain but lacks an N-terminus FERM regulatory sequence. In oligodendrocytes, Ermin was shown to promote 2’,3’-cyclic nucleotide 3’-phospho-diesterase (CNPase) trafficking^4^. Because of the putative interaction of CNPase with microtubules and the role of actin rearrangement in myelinogenesis8, Ermin was suggested to (1) provide links between the actin-based microfilaments and tubulin-based microtubules, and (2) play a role as an interaction hub in the rearrangement of cytoskeletal proteins during myelin wrapping and compaction. Furthermore, *in vitro* crystallography studies suggest that Ermin has a disordered structure, a feature it shares with other cytoskeletal and myelin specific proteins such as myelin basic protein (MBP)^9^. Notably, myelin compaction processes, including MBP polymerization, are thought to begin at the outer layer of myelin sheaths where Ermin is localized^4,10^, supporting a possible role for Ermin in myelin compaction.

Ermin’s expression follows the progression of myelination, appearing in a caudal-to-rostral and ventral-to-dorsal manner, and is found to lag behind MBP by 2-3 days in the postnatal period^4^. Based on its spatial and temporal expression patterns as well as its role in actin binding and remodeling, Ermin has been suggested to play a role in the maintenance of the myelin sheath^5^, although this has not been directly examined in vivo. Furthermore, changes in Ermin expression have been linked to several disorders such as epilepsy^11^, schizophrenia^11,12^, and autism^13^, although no disease-causing mutations in *ERMN* nor a pathological role for it have been reported to date.

Here, we wondered whether loss of Ermin compromised the structural and functional integrity of myelin sheaths. We found that Ermin deficiency led to structural myelin abnormalities and impaired nerve conduction. These deficits were coupled with progressive neurological phenotypes, white matter inflammation and increased susceptibility to an immune-mediated demyelinating insult. Our findings demonstrate an essential role for Ermin in maintaining the integrity of CNS myelin and the pathological consequences of its loss. Furthermore, we have identified a germline inactivating mutation in *ERMN* in a multi-incidence multiple sclerosis family. The etiology of inflammatory white matter disorders such as multiple sclerosis is multi-factorial and associated with significant clinical and pathological heterogeneity^14,15^. While ample evidence supports “outside-in” etiologies in which myelin sheath pathology and degeneration are caused by processes that originate outside oligodendrocytes, an “inside-out” pathogenic process has been more challenging to prove. Given its near exclusive expression in myelinating oligodendrocytes, Ermin deficiency represents a compelling model of “inside-out” inflammatory dysmyelination.

## RESULTS

### Generation of Ermin knockout mice

To investigate the physiological functions of Ermin, we created an Ermin knockout (KO) mouse model using the CRISPR/Cas9 technology^16^. Briefly, Cas9 mRNA and two different guide RNAs (gRNAs) targeting the promoter and exon 1 region of *Ermn* were co-injected into zygotes with a C57BL/6 background (Fig. 1A). As a result, the promoter region, 5’ UTR and 122 nucleotides of exon 1 were deleted. The deletion was confirmed by genomic PCR (Fig. 1B). Immunoblot analysis of brain lysates confirmed ~50% reduction in heterozygotes (HTZ) and complete loss of expression in Ermin knockout (KO) mice (Fig. 1C and D).Knockout mice were born at Mendelian ratios and were viable.

**Figure 1.**
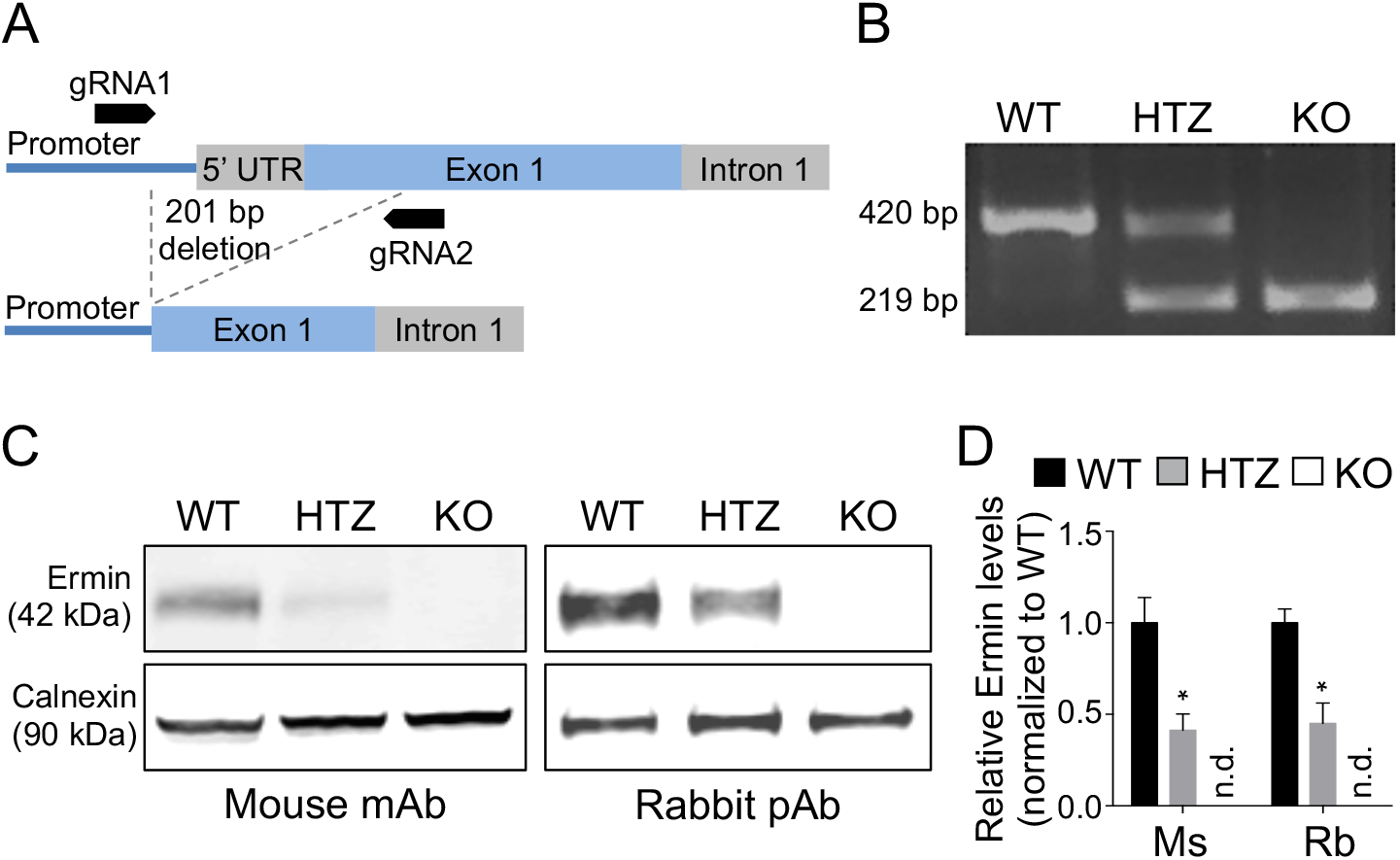
Generation of Ermin knockout mice. (A) Schematic depicting the location of CRISPR/Cas9 gRNA1 and gRNA2 used to generate a 201bp deletion in the *Ermn* gene. (B) Genomic PCR on wild-type (WT), heterozygous (HTZ) and homozygous (KO) mice confirming germline transmission of a 201bp deletion in *Ermn*. (C) Immunoblotting using two different antibodies on whole brain lysates of 3 months old mice confirmed reduction of Ermin protein in HTZ mice and complete absence in KO mice. mAb = monoclonal antibody, pAb = polyclonal antibody. (D) Ermin protein levels are reduced by ~50% in HTZ and absent in KO mice. Ermin levels relative to respective Calnexin loading controls were normalized to WT mice. Values shown as mean±SEM, n = 6 per genotype, * P<0.05 compared to WT was determined by one-way ANOVA with Tukey’s post-hoc test. n.d.=not detected, Ms=mouse antibody, Rb=rabbit antibody.

### Loss of Ermin leads to compromised myelin and axonal degeneration

To assess the impact of loss of Ermin on myelin microstructures, we used transmission electron microscopy to visualize myelinated fibres in the corpus callosum (CC), the largest white matter structure in the brain, in 5 months old mice (Fig. 2A). We evaluated the g-ratio, calculated as the ratio of axon diameter (axon calibre) to myelinated fiber diameter, as a measure of myelin sheath thickness. Plotting g-ratios against axonal diameters demonstrated that g-ratios of larger calibre axons were lower in Ermin KO mice compared with WT littermates (Fig. 2B). In line with these findings, the mean g-ratio of myelinated axons was lower for Ermin KO mice compared to WT, and the cumulative frequency distributions of g-ratios demonstrated a shift to the left in the population, suggesting a thickening of myelin sheaths in Ermin KO mice (Fig. 2B). Furthermore, periodicity, a measure of myelin compaction calculated as the mean distance between two major dense lines, was significantly increased in Ermin KO mice (Fig. 2C and D), suggesting that loss of Ermin leads to de-compacted myelin sheaths in the CC of Ermin KO mice compared with WT mice (Fig. 2A-D). Thus, the increase in thickness of myelin sheaths in Ermin-deficient mice is likely related to its de-compaction in these mice.

**Figure 2.**
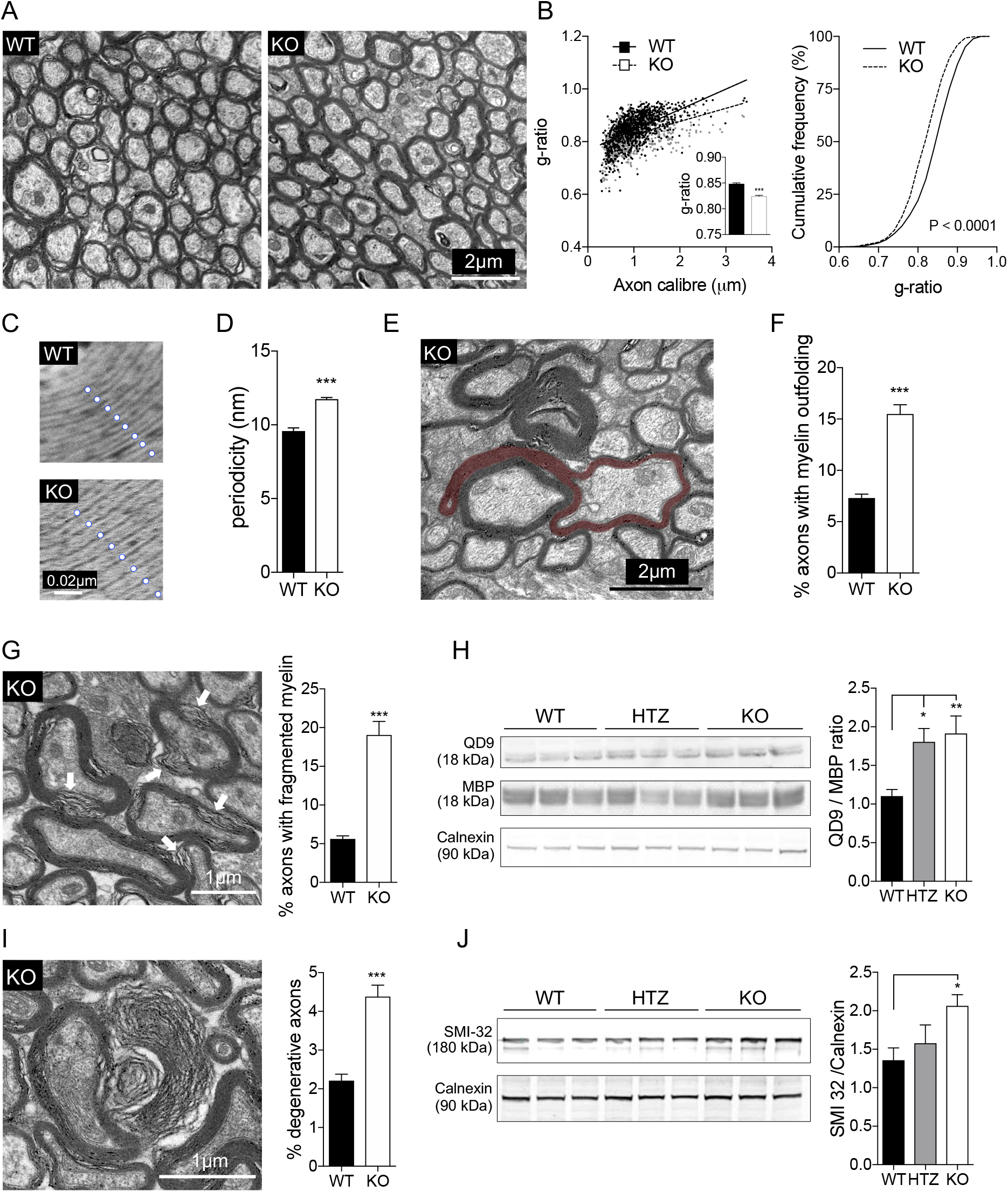
Loss of Ermin results in de-compacted myelin sheath, myelin outfolding and fragmentation, and axonal pathology. Ultrastructural electron microscopy (EM) analysis of myelin was perform on 5 months old mice. (A) Representative EM images of myelinated axons in the corpus callosum in WT and KO mice. (B-D) Ermin KO mice exhibit lower g-ratios (B) and higher periodicity (C and D), suggesting less compact myelin sheath. n=3 per genotype; ~300 axons quantified per animal for g-ratio calculation. Data show means ± SEM; *** P<0.001; two-tailed unpaired Student’s test. (E) Representative EM image of myelin outfolding in the corpus callosum of KO mice. (F) Quantification reveals increased myelin outfolding in Ermin KO mice. Data shown as mean ± SEM; *** P<0.001 by two-tailed unpaired Student’s test. (G) Representative EM image and quantification of axons with fragmented myelin in the corpus callosum of KO mice. Data show means ± SEM; *** P<0.001; two-tailed unpaired Student’s test. (H) Immunoblot showing QD9 and corresponding MBP band, and Calnexin as loading control. Quantification of MBP/QD9 ratio, a measure of myelin damage and fragmentation, at 3 months of age: n = 9/genotype. Data represent means ± SEM; one-way ANOVA with Tukey’s post-hoc test. * P<0.05, ** P<0.001. (I) Representative EM image and quantification of degenerating/ed axons in the corpus callosum of KO mice. Data show means ± SEM; *** P<0.001 by two-tailed unpaired Student’s test. (J) Immunoblot of SMI-32, a marker of axonal damage, and Calnexin as loading control. Quantification of normalized SMI-32 level at 3 months of age: n = 9/genotype. Data represent means ± SEM; one-way ANOVA followed by Tukey’s test. WT vs. KO, * P<0.05.

We next analysed the number of myelinated axons with morphological abnormalities in the CC of Ermin KO mice. Comparing Ermin KO mice to their WT littermates, we observed a significant increase in the percentage of axons with myelin outfolding, a phenotype seen in several myelin gene mutant mice17 (Fig. 2E and F). Similarly, the percentage of axons with fragmented myelin (Fig. 2G) and degenerative phenotype (Fig. 2I) was increased in Ermin KO mice compared with WT mice.

We further investigated whether QD9/MBP ratio was altered. QD9 is an antibody shown to detect an epitope of MBP primarily accessible under conditions of myelin fragmentation or damage^18,19^. We found a significant increase in the QD9/MBP ratio in the brain of 3 months old Ermin KO mice compared with WT (Fig. 2H), indicating compromised myelin sheath integrity in Ermin-deficient mice.

Given the increase in axons with degenerating/ed phenotype, we assessed the levels of SMI32, a measure of the dephosphorylated form of neurofilament proteins and a marker of axonal damage^20^. SMI32 levels were significantly increased in the brain of 3 months old Ermin KO mice, supporting the presence of increased axonal pathology (Fig. 2J). Altogether, our results suggest that loss of Ermin leads to compromised myelin and axonal degeneration.

### Ermin deficiency leads to slower conduction velocity

Facilitation of conduction is one of the primary functions of myelin sheaths. In order to evaluate possible consequences of the compromised myelin sheaths in Ermin KO mice, we recorded the compound action potential (CAP) across the CC of Ermin KO and WT mice at 3 months of age by electrophysiological analysis (Fig. 3A). Quantification of the average stimulus-response time (Fig. 3B) revealed that the conduction velocity was delayed in myelinated fibers of N1 component (Fig. 3C), but not in unmyelinated fibers (N2) (Fig. 3D), in the Ermin KO mice compared with WT mice. Interestingly, myelinated fibers of Ermin Heterozygote (HTZ) mice behaved similarly to those of Ermin KO mice (Fig. 3C). There were no alterations in the amplitudes, area and duration of peaks in CAPs across the CC in Ermin KO mice (Fig. S1). Altogether, these results suggest that loss of Ermin compromises the conductivity of myelinated, but not unmyelinated, fibers.

**Figure 3.**
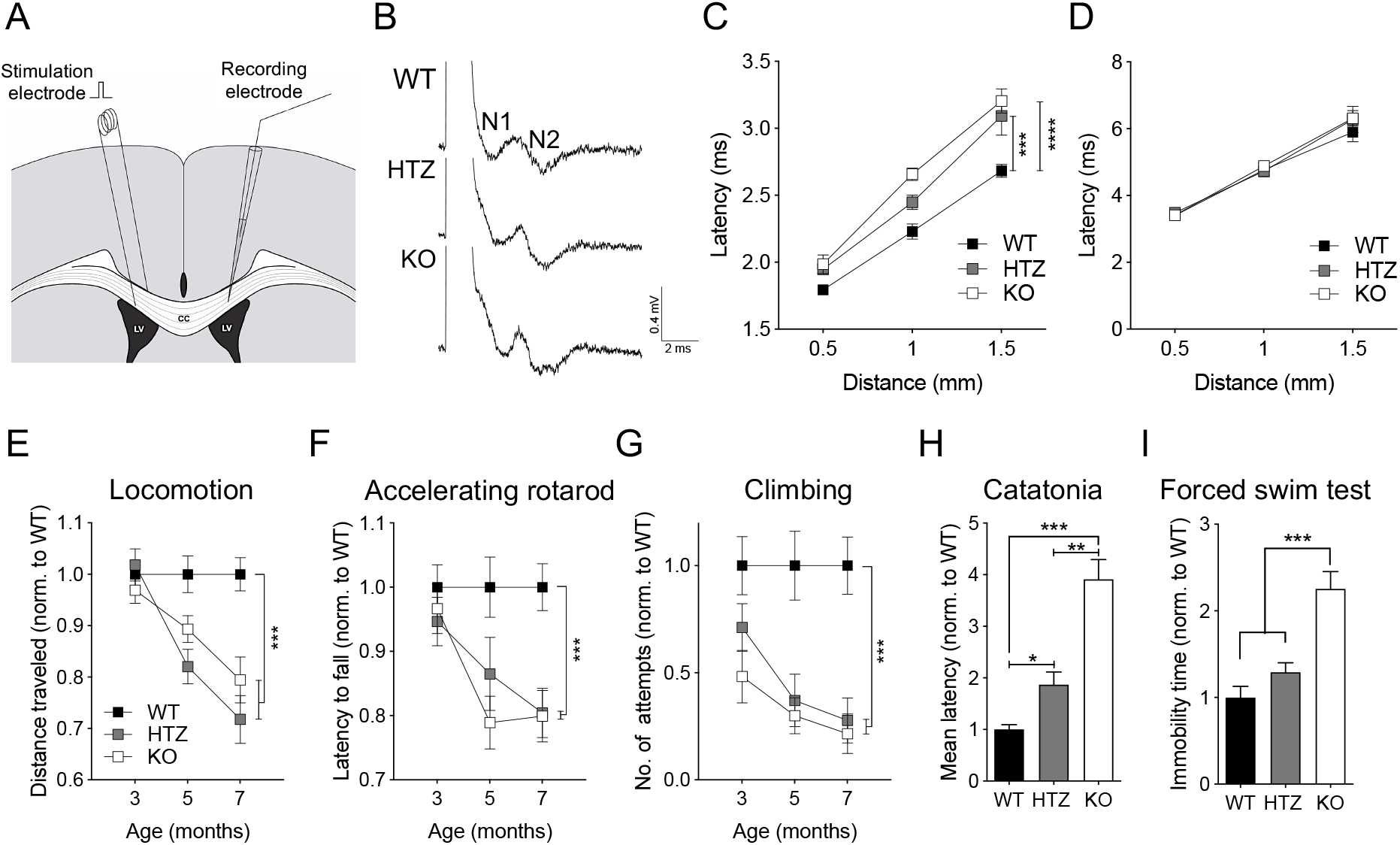
Reduced conduction velocity and progressive neurological deficit in Ermin KO mice. (A) Schematic showing the placement of stimulation and recording electrodes for measurement of compound action potentials (CAPs) in the corpus callosum, (B) representative CAP traces evoked with 5 mA stimuli. (C,D) Conduction was delayed in (C) myelinated fibers (N1), but not (D) unmyelinated fibers (N2) for KO and HTZ mice compared to WT mice. Each plot represents mean ± SEM,n=12 slices from 4 mice per genotypes, *** P<0.001 by two way ANOVA with Tukey’s post-hoc test. (E-G) Ermin-deficient mice exhibited progressive motor deficits based on (E) spontaneous activity, (F) accelerating rotarod performance, and (G) climbing attempts. (H) Ermin KO mice showed catatonic phenotype in the bar test and (I) increased immobility in the Porsolt FST of depressive behaviour at 7 months of age. Data represent means ± SEM; N = 17-25 per genotype; *P<0.05, **P<0.01 ***P<0.001 by (H) Kruskal-Wallis test, and (I) one-way ANOVA or (E-G) two-way ANOVA followed by Tukey’s post-hoc test.

### Progressive neurological deficits develop in Ermin-deficient mice

In light of the myelin structural and functional (conduction) abnormalities, we hypothesized that Ermin deficiency may result in neurological phenotypes. To test this hypothesis, we evaluated mice at 3, 5 and 7 months of age using a battery of behavioural tests. Ermin KO mice exhibited deficits in motor function such as as deficits in spontaneous activity (decrease in distance travelled) (Fig. 3E), accelerating rotarod performance (Fig. 3F), climbing (Fig. 3G). Ermin KO mice also displayed catatonia (Fig. 3H), a phenomenon observed in a number of myelinopathies^21^, as well as depressive-like behaviour in the Porsolt forced swim test (FST) (Fig. 3I). Other behavioural results are summarized in (Fig. S2).

### Transcriptional analysis points to white matter inflammation in Ermin-deficient mice

To gain insights into the processes impacted by the loss of Ermin, we performed RNA-seq analysis on corpora callosal (CC) isolated from 8 months old WT, Ermin HTZ and KO mice (deposited in SRA under accession number PRJNA612161). Principle component reveals general separation of WT samples from the Ermin HTZ and KO samples which showed relatively tighter clustering (Fig. 4A). We identified 579 differentially expressed genes (DEGs; false discovery rate (FDR)<10%) between WT and HTZ, of which 331 genes were upregulated and 248 were downregulated in HTZ. Furthermore, there were 828 differentially expressed genes identified between WT and KO, of which 556 genes were upregulated and 272 were downregulated in KO. Comparison of the differentially expressed genes (DEGs, relative to WT) showed more DEGs in Ermin KO mice compared with HTZ mice (Fig. 4B; Table S1 and S2), suggesting a greater impact of Ermin deficiency in KO mice compared with the HTZ mice. Venn analysis of DEGs showed the transcriptional changes in HTZ mice overlap with those seen in KO mice (Fig. 4C). Functional annotation of the DEGs for Ermin KO mice revealed an enrichment for axonal genes as well as genes associated with myelin sheath and myelination amongst the downregulated DEGs (Fig. 4D; Table S3). Amongst upregulated genes, there was an enrichment of genes associated with immune and inflammatory responses (Fig. 4D; Table S4). A heatmap of representative inflammation-related genes such as *Nfkb1, Itgax*, and *Lcn2* upregulated in Ermin KO mice is shown in Fig. 4E. Overall, the differential gene expression analysis points to inflammatory activation in the corpora callosa of Ermin-deficient mice, amongst other changes related to axonal and myelination processes.

**Figure 4.**
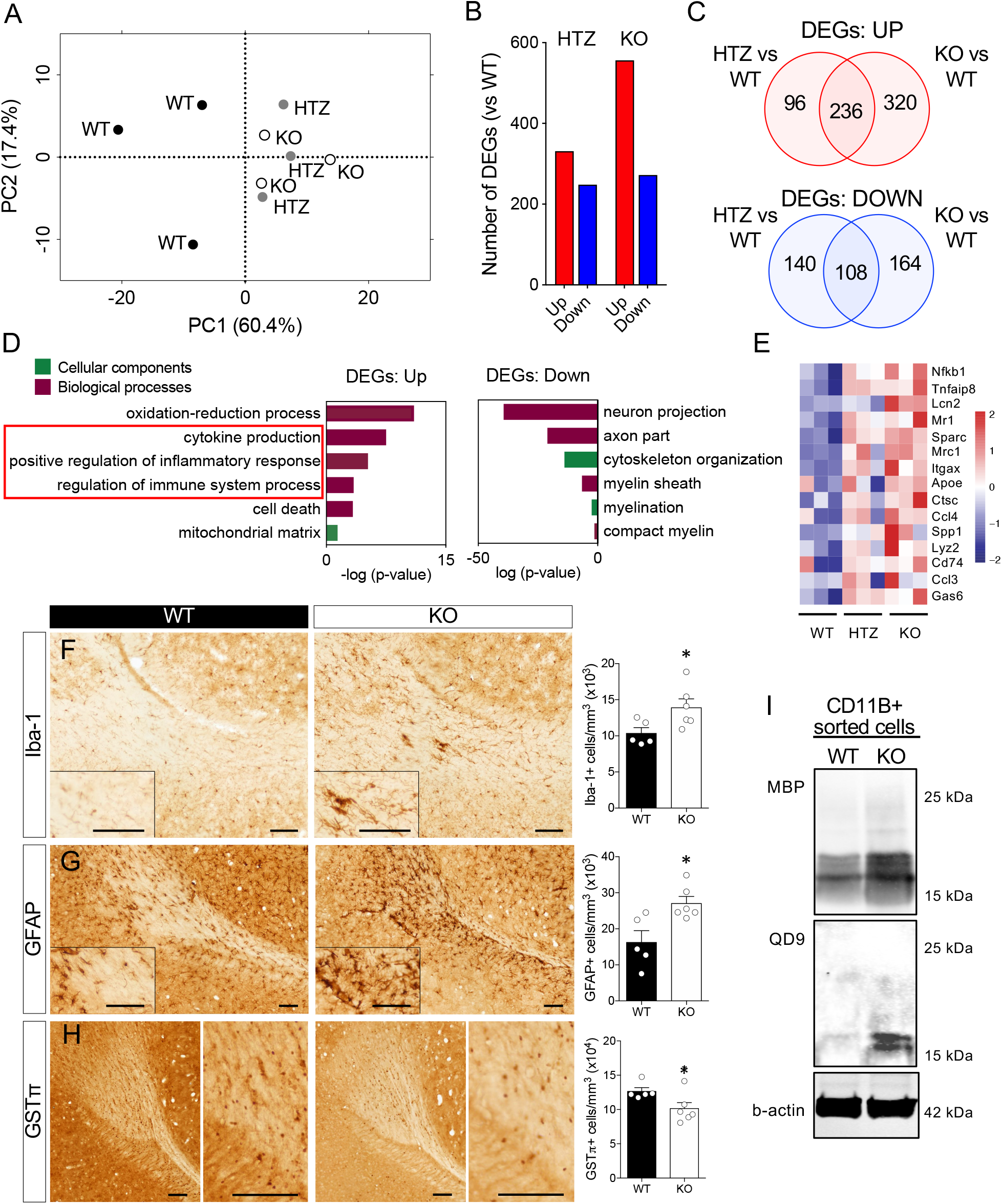
Loss of Ermin leads to inflammatory activation and microgliosis. (A-E) Transcriptional analysis of corpora callosa of Ermin-deficient mice at 8 months of age. (A) PCA plot showing clustering of WTs (n=3), HTZ (n=3) and KO (n=3) mice, (B) The number of deferentially expressed genes (DEGs), both upregulated (UP) and downregulated (DOWN), in HTZ and KO mice compared with WT (FDR<5%). (C) Venn analysis showed a substantial overlap of the UP and DOWN genes between HTZ and KO mice. (D) GO analysis of DEGs between KO and WT mice showed enrichment for terms related to inflammatory processes amongst the upregulated DEGs and myelin-related processes in the downregulated DEGs. (E) Heat map of mean gene expression for selected immune- and myelination-related in WT, HTZ and KO mice. (F-H) Immunohistological analysis shows microgliosis (Iba-1 + counts) (F), astrogliosis (GFAP+ counts) (G), and loss of mature oligodendrocytes (GSTπ + counts) in the corpus callosum Ermin KO mice at 8 months of age. Data show means ± SEM; * P<0.05; two-tailed unpaired Student’s test. (I) Immunoblot analysis of CD11b+ (microglia/macrophages) isolated from brains of 12 months old male mice shows increased levels of phagocytosed MBP and fragmented MBP (QD9).

### Ermin deficiency triggers microgliosis, astrogliosis, and loss of mature oligodendrocytes

To investigate inflammatory and oligodendrocyte changes suggested by the transcriptional analysis, we performed stereological analysis of microglia (IBA-1^+^), astrocytes (GFAP^+^) and mature oligodendrocytes (GST*π*^+^) in CC of 8 months old Ermin KO mice. We observed a significant increase in the total number of microglia (Fig. 4F) and astrocytes (Fig. 4G), and a significant decrease in the total number of mature oligodendrocytes (Fig. 4H) in KO mice compared with WT. These results indicate that loss of Ermin causes increased white matter inflammation and loss of mature myelinating oligodendrocytes.

Clearance of myelin debris is a key function of microglia^22^. To investigate whether the microgliosis observed may be due, at least in part, to an increased demand for clearance of myelin debris in Ermin KO mice, we isolated brain CD11b^+^ cells (microglia/macrophages) and assessed the levels of MBP and fragmented MBP (QD9) they contained. Increased levels of MBP and fragmented MBP were detected in CD11b^+^ cells isolated from Ermin KO cells (Fig. 4I), confirming a higher level of phagocytosed total and damaged myelin in Ermin KO mice.

### Increased susceptibility to inflammatory demyelinating insult in Ermin knockout mice

Immune and inflammatory activation is a feature of several demyelinating white matter disorders. Given the compromise in myelin sheaths and the heightened inflammatory milieu in brains of Ermin KO mice, we sought to examine the impact of Ermin deficiency on the susceptibility to demyelination insult using the widely-used experimental autoimmune encephalomyelitis (EAE) model (Fig. 5A)^23^. EAE-induced pathology is characterised by inflammatory demyelination and neurological phenotypes. In 9 weeks old Ermin KO mice, we observed significantly earlier onset of phenotypic (neurological) deficits compared with WT mice (Fig. 5B,C). Furthermore, a smaller proportion of Ermin KO mice were symptom-free post-induction compared with WT mice (Fig. 5D).

**Figure 5.**
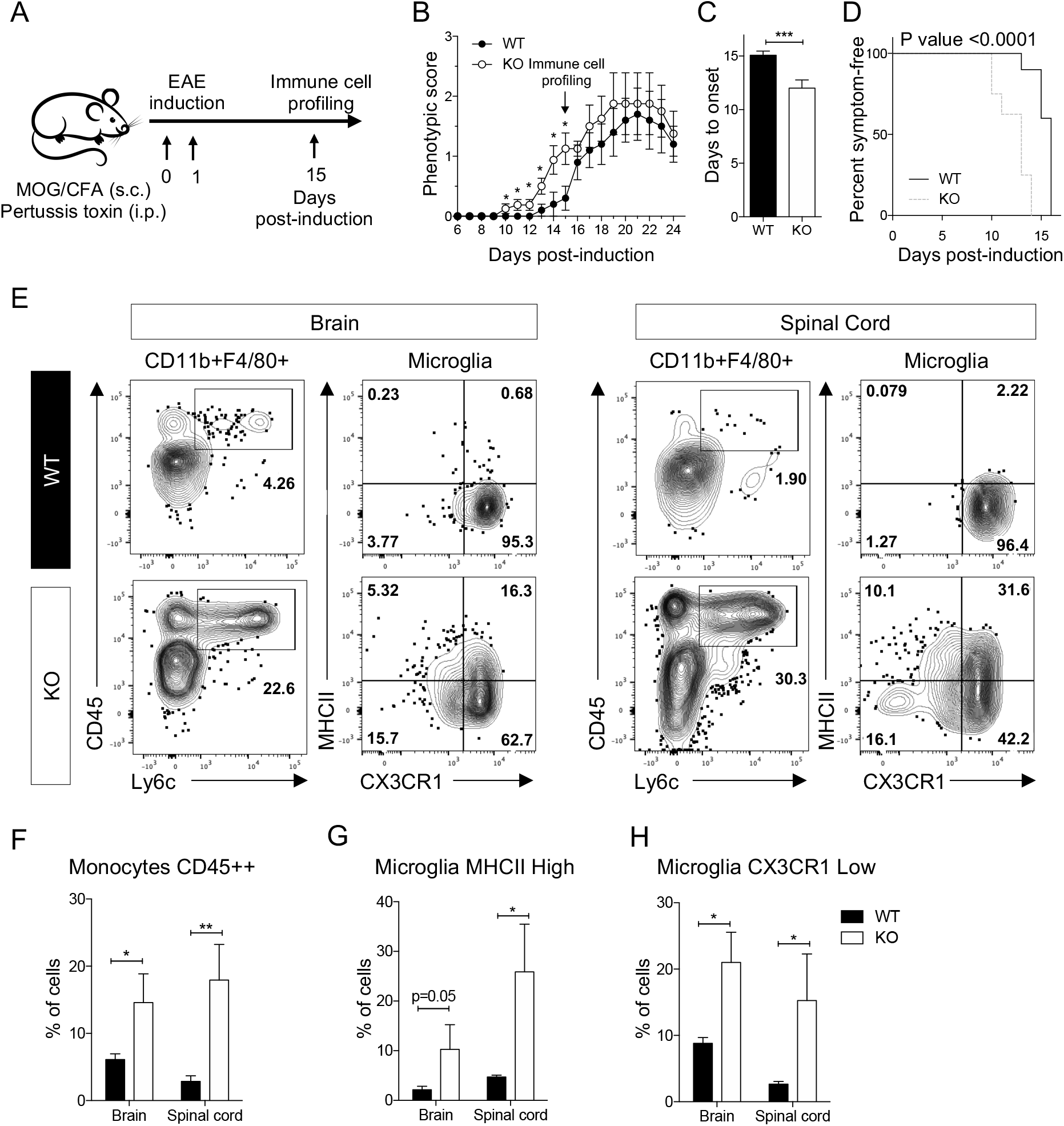
Loss of Ermin increases susceptibility to demyelination insult in the EAE model. (A) Schematic representation of the EAE experiment performed on 9 week old female mice. (B) Daily scoring shows earlier onset of phenotypic deficits in Ermin KO mice compared to WT. (C) Onset of phenotypic deficits is significantly earlier in KO mice compared to WT (D) Analysis of symptom-free mice post-induction shows enhanced susceptibility to EAE in the KO mice (P<0.0001). Graph represents a Kaplan-Meier curve with significance calculated using the log-rank test. (E) Flow cytometry analysis of CD11b^+^ enriched cells isolated from the brains and spinal cords of WT and Ermin KO mice at day 15 of post-induction. The graphs show representative dot plots and related quantification of (F) Monocytes CD45++, and (G,H) Microglia based on (G) MHCII ^*High*^ and (H) CX3CR1 ^*Low*^ levels. Data show means ± SEM; * P<0.05; ** P<0.01; *** P<0.001; two-tailed unpaired Student’s test.

Given such difference in phenotypic onset, we examined the cellular immune response in Ermin KO and WT mice. Flow cytometry analysis demonstrated a massive infiltration of monocytes (CD45^*High*^Ly6C^+^) as well as non conventional monocytes (CD45^*High*^Ly6C^-^) in brain and spinal cord of Ermin KO mice compared with WT (Fig. 5E,F). Microglia constitute the major central nervous system macrophage population and has been shown to play a crucial role in the clearing of myelin in the brain^22^. In fact, we also observed higher levels of fragmented-myelin inside microglia in Ermin KO compared to WT, as well (Fig. 4I). In Ermin KO mice, microglia acquire higher expression levels of MHCII (MHCII^*High*^) (Fig. 5G). MHCII has been described to be associated to microglia activation status^24^. In Ermin KO mice, microglia also down regulates more the fractalkine receptor, CX3CR1 (CX3CR1^Low^) compared with WT mice (Fig. 5H). Down regulation of CX3CR1 has been shown to be associated with the emergence of disease-associated microglia (DAMs)^25^. These data suggest that Ermin KO mice have an increased susceptibility to inflammatory demyelination associated with higher monocyte infiltration, microglia activation, and DAM microglia (CX3CR1^*Low*^) in brain and spinal cord.

### Discovery of mutations in *ERMN* in a multi-incident multiple sclerosis family

Having established the pathological and functional consequences of Ermin loss in mice, we next sought to investigate whether Ermin deficiency could have similar effects in humans. Using GeneMatcher^26^, an online tool that matches researchers with an interest in a common gene, we identified an Iranian family segregating a rare inactivating germline mutation in *ERMN* in which multiple members have been diagnosed with multiple sclerosis (MS). This multi-incident MS family consists of 4 individuals over two generations, two of whom were diagnosed with MS based on the McDonald criteria27 (Fig. 6A,B).

**Figure 6.**
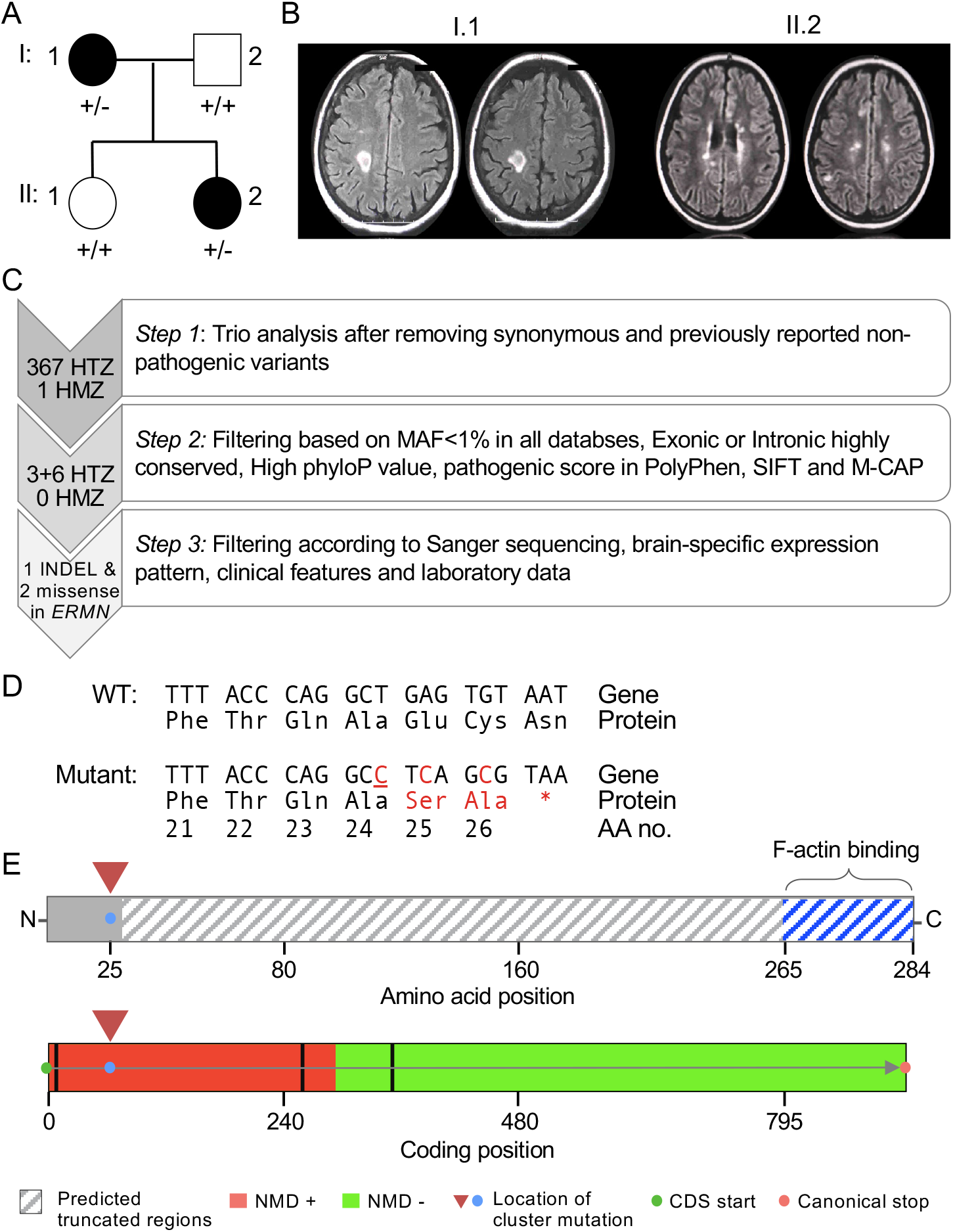
Discovery of novel cluster *ERMN* mutations in multi-incident mutiple sclerosis family. (A) Pedigree of the multi-incidence MS family. Patients diagnosed with MS have black-filled symbols. (B) T2-FLAIR MRI images of the affected mother (I.1) and daugher (II.2) with demyelinating plaques. The mother (I.1) had plaques in the subcortical and periventricular white matter; images show tumefective plaque in the right centrum semiovale. The affected daughter (II.2) had plaques in the pons, basal ganglia, corpus callosum; images show paraventricular and centrum semiovale plaques. (C) Criteria used to filter the variants identified by whole exome sequencing. The filtering approach included trio analysis, *in silico* prediction of conservation and pathogenicity, as well as confirmation by Sanger sequencing. Following filtering, a cluster of three mutations (an insertion and two missense variants) in the *ERMN* gene remained. (D) Predicted amino acid sequence of the mutant *ERMN* allele. Underlined nucleotide represents an insertion. (E) Premature stop codon results in the truncation of ~92% of the ERMIN protein 260/284, including the highly conserved C-terminus actin-binding domain (top). Based on the position of the cluster mutation, *in silico* analysis using the NMDEscPredictor algorithm predicts it to cause nonsense-mediated decay (bottom).

Whole exome sequencing (WES) was performed on the affected mother, affected daughter and healthy father. After applying several filters on the analysed trio, 367 HTZ and 1 HMZ variants remained. The identified variants were further filtered as depicted in (Fig. 6C). Of those, 9 HTZ variants passed filtering criteria that include (a) minor allele frequency (MAF) below 1% in public and proprietary databases of variants, (b) exonic or highly conserved intronic variants (PhyloP score), (c) pathogenic score based on SIFT, PolyPhen and M-CAP (Mendelian Clinically Applicable Pathogenicity)28 (Fig. 6C).

Three adjacent mutations were all found in *ERMN*: two missense mutations and one frame-shift mutation were clustered within 5 nucleotides of each other (Fig. 6D). It is of note that such clustering has been reported as a feature of *de novo* mutations^29^. The cluster mutation is predicted to result in a truncated Ermin fragment that is devoid of Ermin’s highly-conserved C-terminus actin binding protein (Fig. 6D). The mutation is also predicted to trigger nonsense-mediated decay based on *in silico* analysis using the NMDEscPredictor algorithm (Fig. 6E)^30^. We cross-referenced our cluster of SNPs with all publicly available databases such as ExAC, ESP6500, UK10K, 1000G, and dbSNP and none were detected (MAF=0). This finding of an inactivating mutation in *ERMN* in patients diagnosed with MS suggests that similar to our observations in Ermin KO mice, Ermin-deficient may be associated with inflammatory white matter disorders in humans.

## DISCUSSION

Ermin, a CNS-specific, actin-binding myelin protein expressed almost exclusively in mature oligodendrocytes, represents less than 1% of the myelin proteome^6^. Despite its minor abundance, we present here several lines of evidence including structural, functional, transcriptional, and genetic analyses that demonstrate an essential role for Ermin in the maintenance and stability of myelin sheaths and the marked pathological consequences of its loss (summarized in Figure 7).

**Figure 7.**
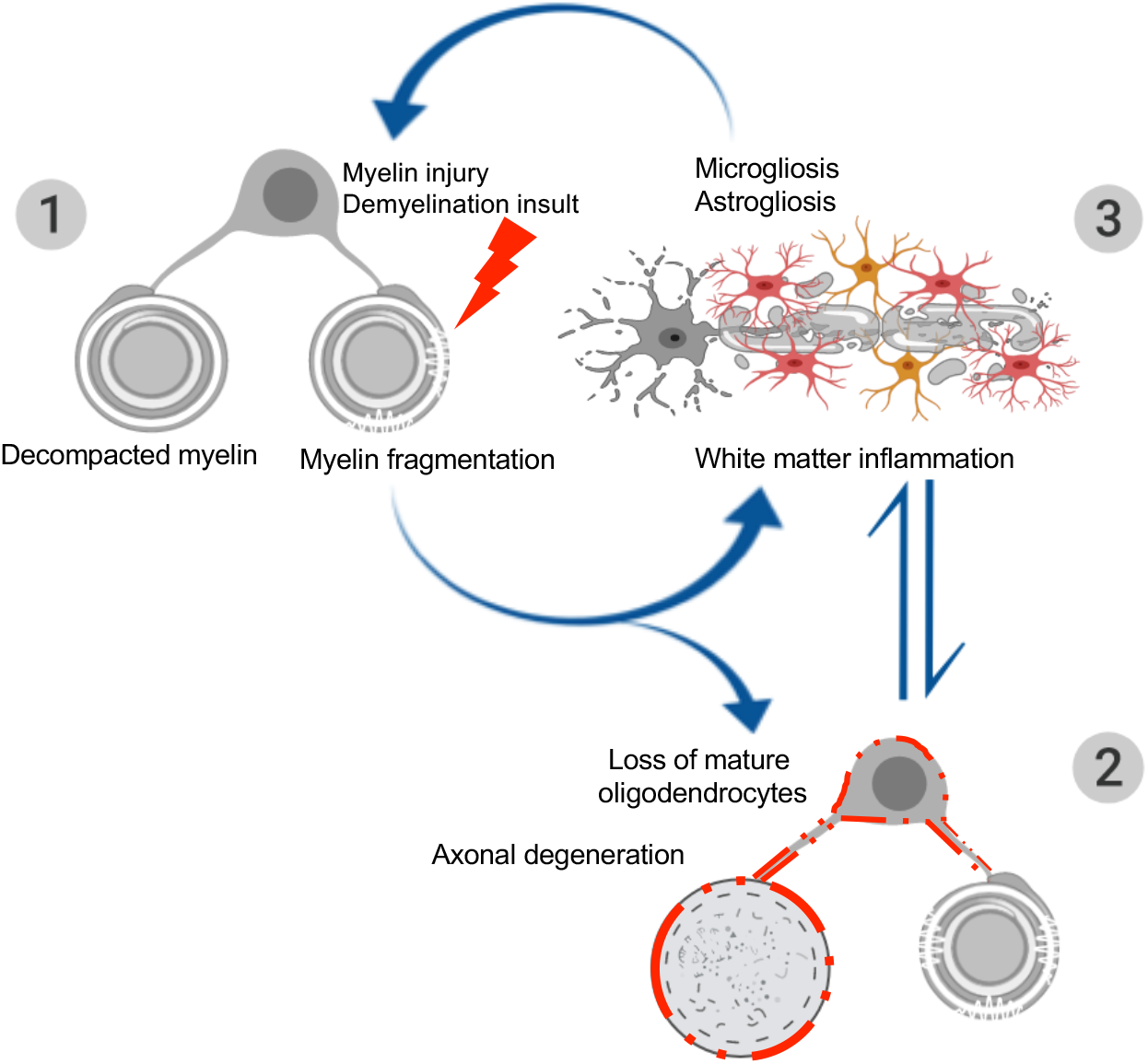
A working model for the sequence of pathological events in Ermin-deficient mice. The pathology in Ermin-deficient mice originates from the myelin sheaths: (1) Loss of Ermin leads to compromised myelin that is de-compacted, outfolded and fragmented; (2) As Ermin-deficient mice age, augmentation of myelin fragmentation and myelin sheath breakdown leads to loss of myelinating oligodendrocytes. Due to compromised myelin sheaths and oligodendrocyte function, loss of trophic support to axons also leads to axonal damage and degeneration; (3) Excessive myelin debris along with oligodendrocyte loss and axonal damage/degeneration triggers white matter inflammation. Activated microglia and astrocytes further exacerbate cell loss. This ensuing inflammation may act as a feed-forward demyelinating insult causing further damage to unhealthy myelin sheaths in Ermin deficient mice.

On the level of myelin ultrastructure, myelinated callosal axons of Ermin-deficient mice exhibited signs of myelin decompaction, fragmentation, and outfolding. Consistent with a compromised myelin sheath, the ratio of QD9/MBP, a measure of myelin damage and fragmentation^19^, was elevated in Ermin-deficient mice. These structural myelin abnormalities were coupled with impaired saltatory conduction as early as three months of age. The reduced velocity of nerve conduction along myelinated callosal fibers in Ermin-deficient mice is likely caused by myelin sheath decompaction and fragmentation which are known to increase axonal capacitance^31^.

In addition to abnormal myelin ultrastructure, signs of degenerating/ed callosal axons were also evident in Ermin-deficient mice. The morphological changes were corroborated by elevated levels of SMI-32, a marker of axonal damage^20, 32^. Since oligodendrocytes and their myelin sheaths play an important role in providing axons with trophic support and neuroprotection^33,34^, the observed manifestations of axonal degeneration provide further evidence of compromised myelin sheath integrity in Ermin-deficient mice.

Ermin deficiency also resulted in a number of neurological deficits, including impaired motor performance, affective phenotypes, and catatonic behaviour. The motor phenotypes were age-dependent, with worsening performance over time suggestive of progressive myelin and axonal pathology in Ermin-deficient mice. Interestingly, catatonia has been shown to be a common neurological manifestation in a number of myelinopathies^35–38^.

Transcriptionally, genes downregulated in the corpus callosum of Ermin-deficient mice showed enrichment for several myelin-related cellular components and biological processes such as “myelin sheath”, “myelination” and “compact myelin”. A number of genes downregulated in Ermin-deficient mice are key components of compacted myelin such as *Mbp, Pllp*, and *Sept8*^17,39,40^, changes that may reflect and/or contribute to the decompaction of myelin sheaths observed in the Ermin-deficient mice. Similarly, and consistent with the signs of axonal pathology observed, gene ontology terms related to axons and neuron death were enriched amongst the genes downregulated in Ermin-deficient mice.

Analysis of genes upregulated in the corpus callosum of Ermin-deficient mice revealed an increase in inflammation and immune-related processes. These include “cytokine production”, ‘‘positive regulation of inflammatory response”, and “regulation of immune system process”. Among these are *Nfkb1*, a master regulator of inflammation^41^, as well as *Mrc1, Itgax* and *Lcn2*, all genes linked to inflammation and innate responses^42–44^. These signs of inflammatory and immune activation were corroborated by histological analyses showing marked microgliosis and astrogliosis in the corpus callosum of aged Ermin-deficient mice.

Microgliosis and astrogliosis are cardinal features of neurodegenerative conditions, including those that involve demyelination and oligodendrocyte loss such as multiple sclerosis^45, 46^. The aetiology and sequence of events leading to these characteristic pathologies and the associated axonal atrophy remain subjects of debate. In the context of multiple sclerosis for example, a dominating view posits that an aberrantly activated immune system initiates a pathological cascade that culminates in oligodendrocyte death and demyelination, paralleled by axonal damage and degeneration. This view represents the initiating trigger as an “outside-in” event^15^. An alternative “inside-out” hypothesis that has been more challenging to prove proposes that the pathology originates within the oligodendrocytes or the myelin sheaths, with the immune and inflammatory activation being a secondary event that propagates the pathogenic cycle^47,48^. Given Ermin’s almost exclusive expression in mature, myelinating oligodendrocytes of the CNS, our study presents compelling evidence in support of the “inside-out” hypothesis. The clinical and pathological heterogeneity of such inflammatory white matter disorders suggest that multiple etiologies, entailing both “outside-in” and “inside-out” pathogenic processes, are likely involved.

How might the microgliosis and inflammatory activation arise in the context of Ermin deficiency? Since microglia play a key role in the removal of myelin debris22, one initiating trigger may be the increased demand for clearance of myelin debris as a result of the instability of myelin sheaths in Ermin-deficient mice. This possibility is supported by the elevated myelin content we detected in brain microglia/macrophages (CD11b^+^) cells from Ermin-deficient mice. Alternatively, the initiating event may be the degeneration of oligodendrocytes as suggested by the decrease in GST*π*-positive cells we observed in the corpus callosum of Ermin-deficient mice. Furthermore, oligodendrocytes are known to express a wide range of immunomodulatory molecules and are thought to be capable of immunomodulation in CNS especially during the initiation of inflammatory responses^49^. Thus, loss or malfunction of oligodendrocytes in Ermin-deficient mice may potentially contribute to the immune activation and microgliosis observed.

Another key feature of a number of demyelinating neurodegenerative disorders such as multiple sclerosis is an increase in the susceptibility to autoimmune-mediated insults. Indeed, many of the therapies currently available for multiple sclerosis act largely through modulation of inflammatory processes^15^. Our findings show that the inflammatory milieu that develops in Ermin-deficient mice is sufficient to increase susceptibility to autoimmune-mediated demyelination. Two further key points can be deduced from these results. First, conditions that compromise the integrity of the myelin sheath are likely to increase susceptibility to immune-mediate demyelination. This is consistent with recent studies in which pharmacologically-induced subtle biochemical changes in myelin sheaths were found to trigger inflammatory demyelination^50^. Second, treatments based solely on the modulation of inflammatory and immune activation in conditions where compromised myelin integrity is the initial trigger are not likely to be effective in the long-term. In this respect, strategies that can improve myelin sheath stability and integrity may be important for effective combinatorial therapeutic interventions.

The possible contribution of impaired Ermin function to human white matter disorders is suggested by our findings of an inactivating mutation in *ERMN* in patients diagnosed with multiple sclerosis. A recent analysis also reported down-regulation of *ERMN* expression in blood samples from patients with multiple sclerosis^51^. While this limited evidence points to dysregulation of *ERMN* in multiple sclerosis, further targeted analyses of Ermin in multiple sclerosis and related conditions is warranted to validate its possible role in susceptibility to demyelinating insults and white matter disorders in humans.

Overall, our findings demonstrate that deficits in factors essential for myelin integrity may contribute to the pathogenesis of inflammatory white matter disorders. Our study is consistent with the notion that low-abundance myelin sheath components such as Ermin may not necessarily be functionally minowr^52^. It further lends support for the “inside-out” theory of inflammatory white matter degenerative disorders, the notion that compromised myelin integrity can trigger inflammation and increase susceptibility to demyelinating insults. Finally, given its near exclusive expression in myelinating oligodendrocytes, the Ermin deficiency model we describe may offer a new paradigm for the development of myelin stability-targeted therapies.

## MATERIALS AND METHODS

### Generation of Ermin knockout mice using CRISPR/Cas9

Ermin knockout (KO) mice were generated on a C57BL/6 background using CRISPR/Cas9. The following guide RNAs that target exon 1 of *Ermn* were used: gRNA_1, “GACGAGTTGGTTTCGAACTGT”and gRNA_2, “GCATACTACAAGGTTGAAC”). Ermin heterozygous (HTZ) mice were bred together to generate Ermin KO mice. Mice were housed with littermates of mixed genotype (2-5 per cage) in individually ventilated cages and kept on a room with 12-h light/dark cycle (lights on at 09:00). Water and food (Altromin 1324 irradiated modified 18% Protein and 6% Fat) were available in cages through study. Mice were maintained under standard conditions and all animal procedures were performed with the approval of the Institutional Animal Care and Use Committee (IACUC # 161186) at Biological Resource Centre (BRC), A*STAR and by their approved guidelines.

### Genotyping

Genomic DNA was extracted from tail tissue of mice using the DNeasy Tissue kit (Qiagen). To visualize the successful deletion of 201 nucleotides of *Ermn* gene in KO mice, the PCR products were run on a 3% agarose gel with SYBR Safe DNA gel stain (Invitrogen). The genotyping primers are: 5’ CCGGGCTGGTTACCAAACT 3’ (forward) and 5’ GATCTTCTCATCTTCAGGCCCTT 3’ (reverse)

### Transmission electron microscopy (TEM)

Mice (n=3 per group) were transcardially perfused with freshly prepared 2.5% glutaraldehyde and 2.5% paraformaldehyde (PFA) in phosphate buffered saline (PBS). Whole brains were post-fixed overnight at 4°C in the same fixative solution, washed in PBS, and stored in 5% sucrose and 0.1% sodium azide at 4°C. Coronal slices at the level of Bregma −1 mm were obtained consistently from one hemisphere using a stainless steel mouse brain slicer matrix (Agnthos), followed by micro-dissection of the anterior region of the corpus callosum (~Bregma 1.10-0.50 mm, according to the Mouse Brain Atlas,53) and washed in PBS. Tissues were then post-fixed in 1% Osmium tetroxide for 1 hour at room temperature (RT), dehydrated through an ascending ethanol series (25% - 100%) and 100% acetone wash, and infiltrated with 1:1 acetone:resin for 30 min at RT and 1:6 acetone:resin overnight at RT. Sections were then transferred to 100% resin for embedding and polymerization at (30 min at 40°C, 1 hour at 45°C and 50°C, respectively, and 60°C for 24 hours). Ultrathin sections (90nm) were obtained using Ultracut E with a diamond knife (Diatome, Ultra45, 3mm length) attachment and embedded on a copper grid. TEM imaging was performed on an FEI TECNAI Spirit G2 for g-ratios and a JEM1010 Microscope equipped with bottom-mount SIA model 12C high resolution full-frame CCD camera (16bit, 4K) for periodicity. Over 300 axons and myelin fiber diameters per animal were measured using ImageJ. G-ratios were obtained by calculating the ratio between axon diameter and axon diameter plus the outer most layer of surrounding myelin. G-ratios were calculated within equal axonal distributions. Periodicity were calculated by dividing myelin thickness to number of myelin layers.

### Protein analysis

RIPA buffer (Sigma-Aldrich) with 1mM PMSF (Sigma-Aldrich), 5mM Z-VAD (Promega), 1mM NaVan (Sigma-Aldrich), and 1x Complete Protease Inhibitor Cocktail tablets (Roche) were used to prepare Protein lysate of half brain from mice. For western blotting 30 g of protein lysates were separated on 12% Bis-Tris protein gel (Novex) with 20X MES running buffer (Novex) and transferred on nitrocellulose membranes. Primary antibodies (listed in Table 1) were incubated at 4°C overnight. Secondary antibodies (1:10,000) were used: Alexa-Fluor goat anti-rabbit 800, Alexa-Fluor goat anti-mouse 680, Alexa-Fluor goat anti-rabbit 680 and Alexa-Fluor goat anti-rat 800 (all from Life Technologies). The membrane was imaged using the LiCor Imaging System and Odyssey V3.0 software (LiCor), followed by intensity analysis with imageJ.

**Table 1.**
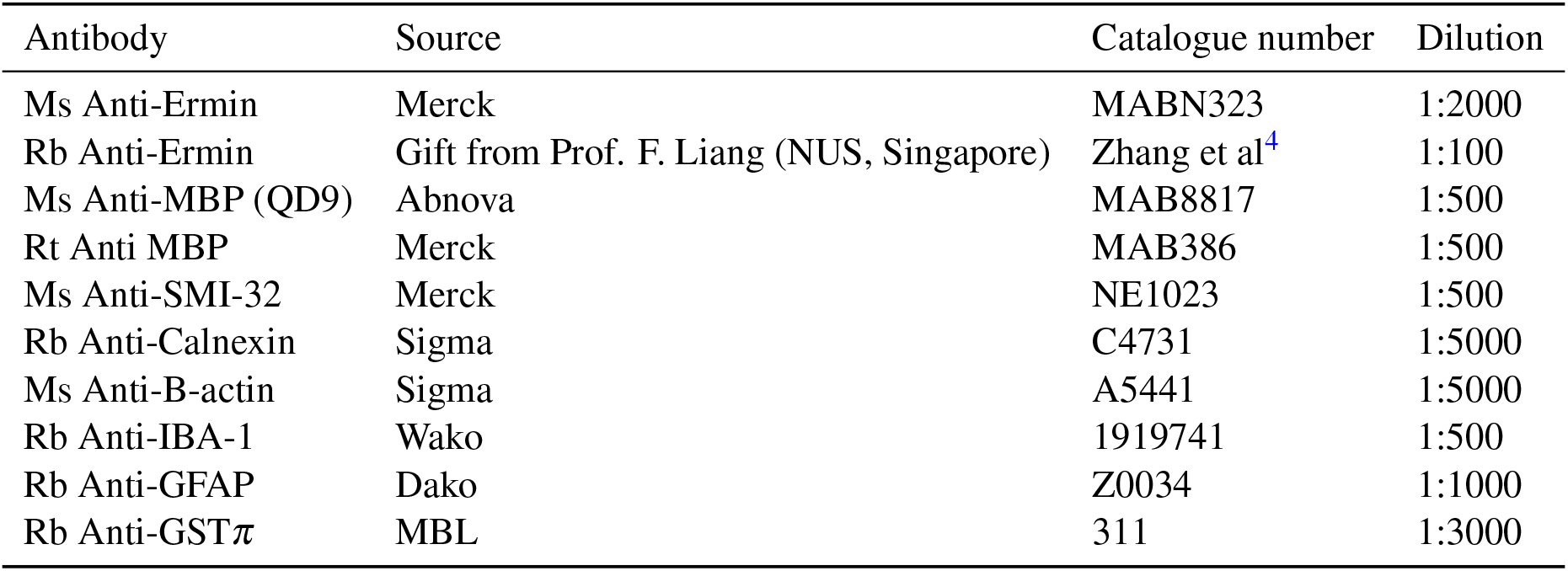
Antibodies used for immunobloting and immunohistochemistry

| Antibody | Source | Catalogue number | Dilution |
| --- | --- | --- | --- |
| Ms Anti-Ermin | Merck | MABN323 | 1:2000 |
| Rb Anti-Ermin | Gift from Prof. F. Liang (NUS, Singapore) | Zhang et al<sup>4</sup> | 1:100 |
| Ms Anti-MBP (QD9) | Abnova | MAB8817 | 1:500 |
| Rt Anti MBP | Merck | MAB386 | 1:500 |
| Ms Anti-SMI-32 | Merck | NE1023 | 1:500 |
| Rb Anti-Calnexin | Sigma | C4731 | 1:5000 |
| Ms Anti-B-actin | Sigma | A5441 | 1:5000 |
| Rb Anti-IBA-1 | Wako | 1919741 | 1:500 |
| Rb Anti-GFAP | Dako | Z0034 | 1:1000 |
| Rb Anti-GSTπ | MBL | 311 | 1:3000 |

### Corpora callosa slice preparation and electrophysiology

Three months old male mice were used to measure compound action potentials (CAPs) in CC. After cervical dislocation, brains were dissected and placed in oxygenated (95% O_2_ + 5% CO_2_) ice-cold sucrose based artificial cerebrospinal fluid (aCSF) solution (206 mM Sucrose, 2 mM KCl, 1.25 mM NaH_2_PO_4_, 26 mM NaHCO_3_, 10 mM Glucose, 2 mM MgCl_2_, 2 mM MgSO_4_, 1 mM CaCl_2_; 305-315 mOsm, pH 7.3 - 7.4). Brains were sliced coronally in sucrose based ACSF using vibratome (Leica VT1200s) with 450 m thickness, and slices were collected approximately from Bregma 1.10 to −0.22 (3 slices per mouse) in Mouse Brain Atlas (Franklin and Paxinos, 3rd ed.). Slices were immediately transferred to a holding chamber filled with pre-warmed (37°C) oxygenated (95% O_2_ + 5% CO_2_) standard aCSF solution (124 mM NaCl, 2.5 mM KCl, 1.2 mM NaH_2_PO_4_, 24 mM NaHCO_3_, 5 mM HEPES, 12.5 mM Glucose, 2 mM MgSO_4_, 2 mM CaCl_2_), and the slices were recovered for 30 minutes at 37°C, afterward kept at room temperature till recording.

Compound action potentials (CAPs) evoked by electrical stimulation were recorded in the corpora callosa at three different distances between stimulation and recording electrodes: 1.5 mm, 1.0 mm and 0.5 mm distance. The recording electrode assembled glass pipette (1-3 M) filled with 3 M NaCl was placed 1 mm away from the midline of brain slice, and stimulation electrode was placed 0.5 mm away from middle line on the opposite side of recording pipette. The recording was conducted with order from 1.5 mm to 0.5 mm by moving in recording electrode. The intensity of the stimulus was increased from 0.5 mA to 5 mA using external stimulator (model 2100; A-M System), and responses were recorded at each step in triplicate. The Multiclamp 700B, Digidata 1550B and pClamp10 software (Molecular Devices) were used for recordings. From acquired data, amplitudes, area of N1 (myelinated) and N2 (non-myelinated) were calculated using Population spike analysis tool in AxoGraph X (AxoGraph Scientific). The area and duration of peaks on 5 mA stimulation at 1 mm distance were used to compare compared between groups.

### Study design of behavioural analysis

All behavioural tests were performed during the dark phase of the reversed light/dark-cycle. Two cohorts of mice were behaviorally tested at 3, 5, and 7 months of age. Mice were evaluated for motor and psychiatric-like behavioural. For all behavioural tests, mice were allowed to acclimatise to the testing room for at least 30 min prior the commencement of the behavioural test. Investigator was blinded for all test sessions.

### Spontaneous locomotor activity test

The spontaneous locomotor activity test is used to measure gross and fine motor movements in mice^54^. The aspects of activity measured are horizontal activity such as distance traveled, and vertical activity such as vertical and jump counts. In order to assess motor function, mice were monitored for 30 minutes using the Med Associates spontaneous activity chambers (27.3 [L] x 27.3 [W] x 20.3 [H] cm) with 16 beams (Med Associate Inc).

### Rotarod test of motor coordination

Motor coordination and balance were evaluated using the accelerating rotarod task performed on a UGO Basile 47600 Rotarod with a rotating rod diameter of 3 cm^55^. Training was performed at three months of age and consisted of three trials (120 s) per day at a fixed-speed of 18 rpm for three consecutive days. Testing was performed at 3, 5, and 7 months of age, and consisted of three trials, spaced 2 hours apart, with a rod acceleration from 5 to 40 rpm within 5 min. Rotarod scores are the average of three trials.

### Climbing test

The climbing test is used to evaluate motor function in mice^56^. Testing was performed at 3, 5, and 7 months of age. Mice were placed at the bottom end of a closed-top metal wire mesh cylinder (10.5 cm of diameter and 15.5 cm high) and were video recorded for 5 min. Number of climbing event were counted manually and blinded. Climbing was determined when all four paws of the mouse were on the walls of the cylinder.

### Elevated plus maze test of anxiety

The Elevated plus maze (EPM) test is classically used to assess anxiety in rodents^57^. The EPM apparatus is a cross-like shape with two open arms perpendicular to two closed arms of equal dimensions. The closed arms are surrounded by three 10-cm high walls. Because mice have an innate fear of high open spaces, they prefer to spend less time in the open arms. Time spent in the open arms is taken as a measure of anxiety-like behavior. Test sessions lasted 5 minutes and time spent in the open arms were recorded using an automated video-based tracking system (Noldus EthoVision 9, Netherlands).

### Porsolt forced swim test (FST) of depression

The Porsolt FST is used to evaluate depressive-like behaviour in mice. The test was performed at seven months of age as previously described^56^. Mice were put in individual cylinders (25 [H] x 19 [W] cm) filled with room temperature water (23-25°C) to a depth of 15 cm for a period of 6 min. The test sessions were recorded and examined blinded. The last 4 min of the test session was scored using a time-sampling technique to rate the predominant behavior including swimming and immobility over 5-sec intervals. Time spent immobile is considered as a measure of depressive-like behaviour.

### Bar test of catatonia

This test was performed as previously described^35,37^. Briefly, the mouse was carried by the tail to a horizontal bar made of stainless steel (12 [L] x 4-5 [H] cm, and 2.5 cm of diameter) and allowed to grasp the bar with both forepaws. Upon grasping the bar with both forepaws, the mouse was moved downwards so that its hind paws had contact with the floor before its tail was released. The time a mouse stood nonmoving with at least one forepaw on the bar and both hind paws on the ground was measured. The test was performed three times for each mouse, and the catatonia score for each mouse was the average of the three trials.

### Brain tissue collection for molecular and sequencing analysis

Corpora callosa form 8-month-old animals were dissected, snap-frozen in liquid nitrogen and stored at −80°C until further use.

### RNA-Seq analysis

RNA was extracted from mouse corpora callosa (CC) using Trizol (Life Technologies) and subsequently a RNeasy plus mini kit (Qiagen) according to the manufacturer’s instructions. Subsequent library preparation and paired-end 150bp sequencing and 15M reads/per sample using HiSeq were performed by Novogene (Hong Kong).

RNA-Seq data analysis was done on the AIR platform (www.transcriptomics.cloud). The trimming is performed with BBduk to remove low-quality bases (minimum quality 25, minimum length 35 bp). High-quality reads were then aligned to the mm10 genome (Ensembl version 89) using STAR 020201 (https://github.com/alexdobin/STAR) and quantified using featureCounts (http://bioinf.wehi.edu.au/featureCounts/), genes expressed at a low level were removed with the package HTSFilter (http://www.bioconductor.org/packages/release/bioc/html/HTSFilter.html), Differential expression of the filtered genes was analyzed by NOIseq method, pairwise. Genes were considered statistically differentially expressed if the corrected P-value (false discovery rate, FDR) was <10%. Enrichment analysis was performed using Hypergeometric Optimization of Motif EnRichment (HOMER)58 on the proportion of GO categories between the differentially expressed genes (DEGs) and the background of expressed genes; GO categories were considered enriched if the FDR of the test was <0.05. An expression heat map was constructed for a pre-selection of genes of interest using the web-based tool Heatmapper (https://software.broadinstitute.org/morpheus/), transforming transcripts per million (TPM) values using a Z-score scaling.

### Immunohistochemistry and stereological measurements

For immunohistochemistry one cohort of mice was used (N = 6-8 per genotype). From all brains, one hemisphere was post-fixed overnight in 4% PFA and switched to 30% sucrose in 1X phosphate buffered saline (PBS) in the following days, and the other was cut via cryostat (Leica CM3050S) into a series of 25m sagittal sections free-floating in 1X PBS with 0.01% sodium azide. For all stainings, sections were washed three times with 1X PBS, and then blocked in 5% normal goat serum (NGS) in PBS containing 0.1% Triton X-100 (TX) for 120 minutes at room temperature (RT). Sections were then incubated with primary antibodies in 1% NGS in PBS-TX overnight at 4°C. The following antibody concentrations were used: rabbit anti-GST*π* at 1:3000 (MBL, #311), rabbit anti-IBA1 at 1:500 (Wako, #01919741) and rabbit anti-GFAP at 1:1000 (Millipore, #OBT0030). Sections were washed three times with PBS and stained using biotinylated anti-rabbit secondary antibody (1:1000, Vector Laboratories) for 90 minutes at RT. Sections were then incubated using ABC Elite Kit (Vector Laboratories) to amplify signal, and then in 3,3’-diaminobenzidine (DAB, Vector Laboratories) for signal detection. For stereological measurements, corpus callosum (CC), fimbria, hippocampus and caudate regions from one hemisphere were traced using the StereoInvestigator software (MBF Bioscience). IBA1 positive cells were counted in all regions while GFAP and GST*π* positive cells were counted in the CC using the Optical Disector probe. The following parameters were used: Iba1 - 100μm x 100μm counting frame, 450μm x 450μm sampling grid; GFAP - 150μm x 150μm counting frame, 300μm x 300μm sampling grid; GST*π* - 50μm x 50μm counting frame, 300μm x 300μm sampling grid. Volume of counted areas was determined using the Cavalieri Estimator probe. Activated microglia and astrocytes were visually identified by increased cell body size and more intense IBA1 or GFAP staining^59,60^. Another significant morphology of activated microglia and astrocytes is contracted ramifications that appear thicker than in their resting state.

### Experimental autoimmune encephalomyelitis (EAE) induction and evaluation

Mice were immunised by Hooke kits (Hooke laboratories, EK-0115, Lawrence, MA, USA) according to the manufacturer’s instructions. Briefly, 0.1 ml MOG35-55/CFA emulsion was injected subcutaneously into both flanks of each mouse (0.2 ml/animal) after anaesthesia. Then, pertussis toxin (0.1 ml/animal/day, i.p.) was injected intraperitoneally (IP) on the same day and 24 hours later.

Two researchers scored the clinical signs of mice daily from day 6 to day 25 post-immunisation using the scale 0 to 5 point EAE scoring system was based on Hooke lab protocol briefly as follows: 0, no clinical sign; 0.5, Tip of tail is limp; 1.0, complete tail paralysis; 1.5, Limp tail and hind leg inhibition; 2.0, Limp tail and weakness of hind legs.; 2.5, unilateral hind limb paralysis; 3, complete hind limb paralysis; 3.5, hind limb paralysis and forelimb weakness; 4.0, complete paralysis (tetraplegia), and 5.0, moribund or dead. The successful induction of EAE in all WT and KO mice was confirmed by this scoring scale. These clinical scoring results were analysed to compare the course of EAE between the two groups.

### Microglia and macrophages enrichment protocol

Brain or spinal cord were harvested and put in 3mL or 1ml solution, respectively, comprised of RPMI, 10%FCS, Collagenase (0.2mg/mL), and DNAse (30*μ*g/mL) using 12 wells plates. Cut in small pieces and incubated at 37°C for 30min. Using 3mL syringes with blunt needle the tissues were dissociated by aspiring and flushing 10 times. Cell suspension was filtered on a cell strainer 70*μ*M and falcon tube 50mL. the cells were washed with 14mL of FACS buffer (PBS+5% BSA+2mM EDTA) and centrifuge 4min at 1600rpm 4°C. Supernatant was removed and cells were resuspended in 1mL FACS buffer + 100*μ*L of microbeads anti-mouse CD11b for brain and spinal cord was resuspended in 200*μ*L + 20*μ*L of microbeads. Cells were incubated for 15min at 4°C. 5mL of FACS buffer was added after incubation and cells were centrifuged at 1600rpm 4min 4°C. Supernatant was removed and 2mL of FACS buffer added. Using the automacs possel function CD11b+ cells were retrieved. CD11b+ cells were centrifuged at 1600rpm 4min 4°C.

### Microglia and macrophage flow cytometry (FACS) staining

Antibodies were diluted in blocking buffer containing FACS buffer+1% rat serum+1% mouse serum. All antibodies were used at 1/200 dilution. 100*μ*l of staining mixture was used per brain and 50uL was used per spinal cord. Cells were incubated for 15min with antibodies mixed at 4°C and then washed with 1mL of FACS buffer and resuspended in 200*μ*L of PBS+DAPI (1*μ*g/mL). The list of FACS antibodies used is listed in Table 2.

**Table 2.**
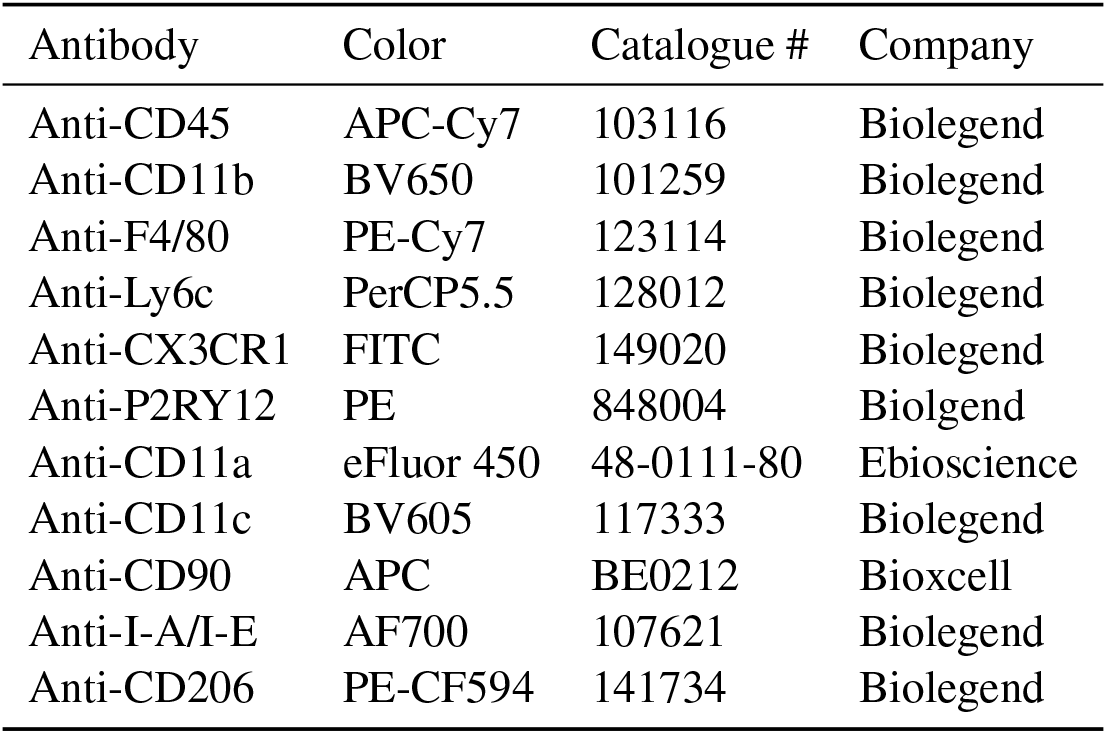
FACS antibodies used

### Discovery of novel mutations in *ERMN* in a multi-incident MS family

Board-certified clinical neurologist evaluated all patients clinically. The entire family was recruited under an Isfahan University of Medical Sciences ethics-approved research protocol with informed consent.

We applied whole exome sequencing (WES) analysis to a multi-incident MS family consisting of 4 individuals over two generations, two of whom were diagnosed with MS For WES, 1*μ*g of gDNA of healthy father, affected mother and affected daughter (1, 2 and 3) used for exome library preparation on an Ion OneTouch System. Amplified samples were sequenced on an Ion Proton instrument (Life Technologies, Carlsbad, CA, USA) using one ION PI chip. Sequence reads were aligned to the human GRCh37/hg19 assembly (UCSC Genome browser). The variants were filtered based on functional prediction scores including SIFT, PolyPhen2, Grantham and M-CAP (Mendelian Clinically Applicable Pathogenicity), as well as PhyloP conservation scores. Variants were further filtered for common single nucleotide polymorphisms (SNPs) using “common and no known medical impacts” database (ftp://ftp.ncbi.nlm.nih.gov/pub/clinvar/vcf_GRCh37/) and the Exome Aggregation Consortium (ftp://ftp.broadinstitute.org/pub/ExAC_release/release0.2/). Variants were next compared to an in-house (IMCB) database of 332 previously sequenced samples, and those that were present in more than 1% of the previously sequenced samples were removed. WES of family members 1,2, and 3 gDNA generated a total of 14.5 Gb with an average read length of 189 bp. An average coverage of 181.9X was achieved across the exome, with 96.89% of the targeted sequences covered at 20X. A total of 67,763 variants were identified across protein-coding exons, untranslated region (UTRs), splice sites and flanking introns. Sanger sequencing was used to confirm that the identified mutation segregates in the family. The primers used for Sanger sequencing were: 5’-ggacaattgtggctcctgtt-3’ (forward) and 5’-ggtgtccagctcatttcgat-3’ (reverse).

### Accession numbers

The RNA-Seq data reported in this paper have been deposited into the Sequence Read Archive under accession number: PRJNA612161.

### Statistical Analysis

Prism 6 (GraphPad Software) was used for all statistical analyses. Student’s t-test was used to calculate statistical significance When only two experimental groups were compared, unless otherwise stated. For multiple comparisons, a one-way or two-way ANOVA was performed followed by a post-hoc Tukey’s multiple comparisons test. For all other experiments. Differences were considered statistically significant when P <0.05.

## AUTHOR CONTRIBUTIONS

AZ and MAP designed research; AZ, MGM, CIR, AS, HGB, HS, NABMY, CFB, LJT, ST, and LD performed research; AZ, AS, HGB, CB, AYJN, SRL, AYJN, and MAP analysed data; BV, VS, BR, SJ, FG provided intellectual input; AZ and MAP wrote the paper with feedback from the other authors; and MAP conceptualised the study.

## ACKNOWLEDGEMENTS

We thank members of the Pouladi and Reversade labs for helpful discussions and comments. We thank Dr Fengyi Liang for the gift of the polyclonal anti-Ermin antibody and helpful discussions. We thank Dr Ehsan Ziaei, Dr Giti Sadeghian, Sahar Dehghan Kelishadi, Shaun Tan and Eri Aung for technical assistance and the A*STAR Microscopy Platform for assistance in sample imaging. The work was partly funded by a Strategic Positioning Fund for Genetic Orphan Diseases (SPF2012/005) and SUREKids (IAF311019) from the Agency for Science Technology and Research (A*STAR, Singapore) to MAP and BR, and a Joint Council Office grant (BMSI/15-800003-SBIC-00E) from A*STAR to SJ.

---

**Figure S1.**
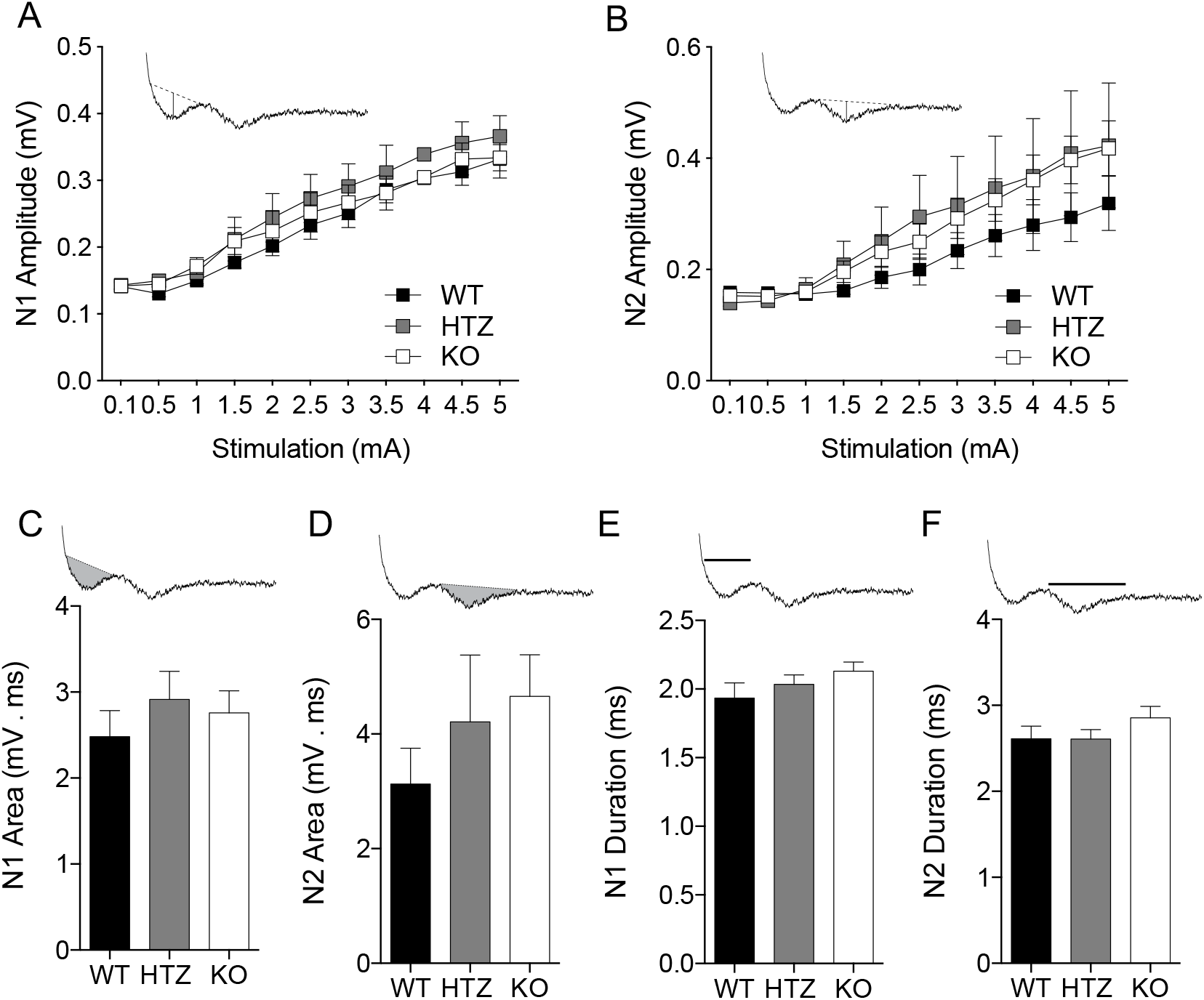
No alteration in amplitudes of CAPs across CC in Ermin KO mice. The amplitude was recorded in 1 mm distance between stimulation and recording electrodes with various stimulation strength (from 0.1 to 5 mA), and the averaged amplitudes of N1 (A) and N2 (B) were plotted against simulations. The areas (N1, C; N2, D) and duration (N1, E; N2, F) were calculated from the traces responded to 5 mA stimulation at 1 mm distance. The traces show the way to measure the area (C, D) and duration (E, F). Data were represented with means ± SEM; 12 brain slices (n) from 4 mice (N) per genotype.

**Figure S2.**
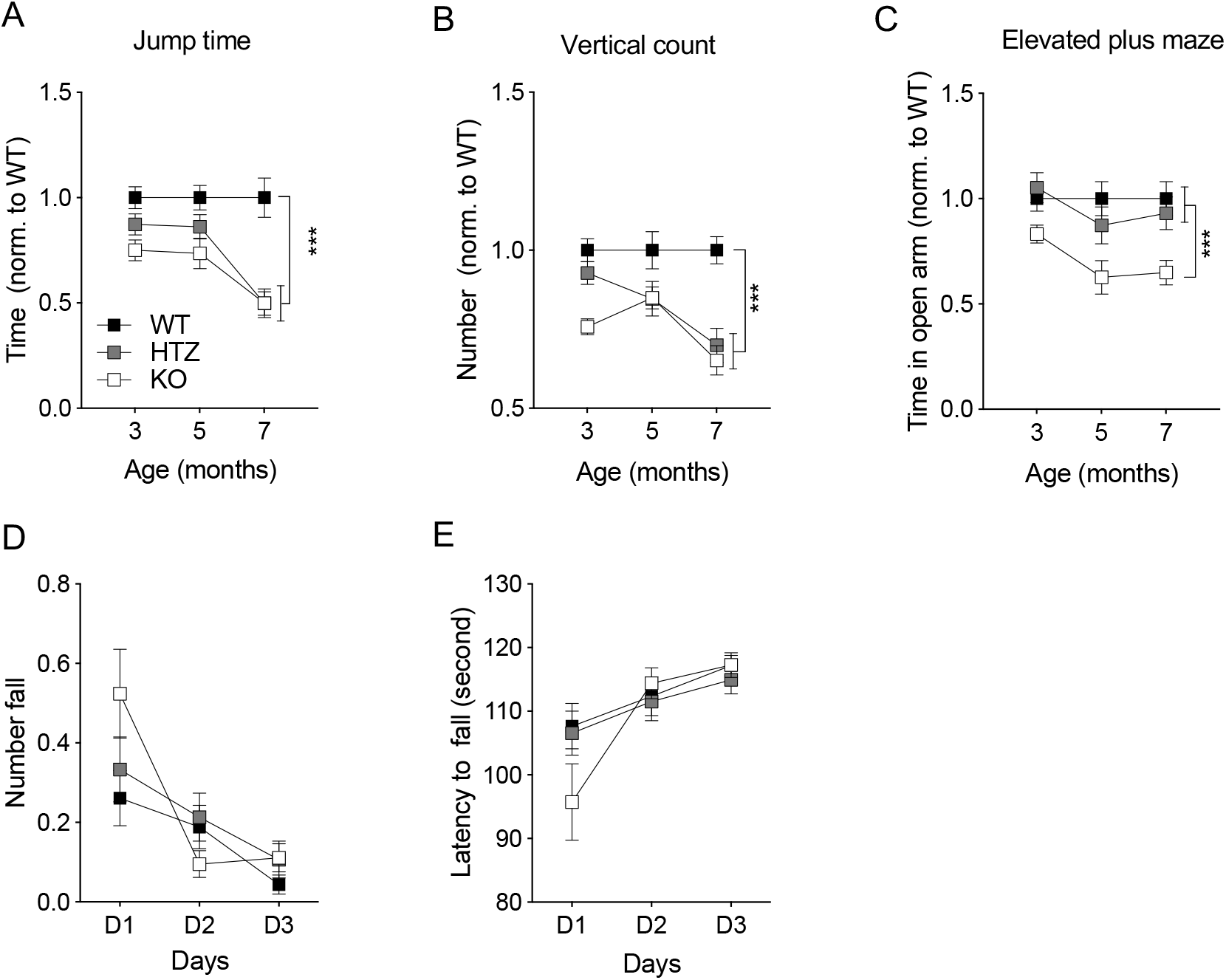
Additional behavioural assessments of the Ermin knockout mice. (A and B) Mice were assessed for motor symptoms. progressive locomotion deficits in the jump time and vertical count in 30 minutes (C). Mice were assessed for anxiety-like symptoms by elevated plus maze. (D and E) Rota road training test in three consecutive days as a cognitive assessing test. N = 17-25 per genotype, littermates, mixed gender. Data show means ± SEM; *** P < 0.001; Two-way ANOVA followed by Tukey’s multiple comparisons test were applied for all behavioral studies.

